# SREBP1 drives KRT80-dependent cytoskeletal changes and invasive behavior in endocrine resistant ERα breast cancer

**DOI:** 10.1101/380634

**Authors:** Ylenia Perone, Aaron J. Farrugia, Alba Rodriguez Meira, Balázs Győrffy, Charlotte Ion, Andrea Uggetti, Darren Patten, Antonios Chronopoulos, Monica Faronato, Sami Shousha, Jenny H Steel, Claire Davies, Naina Patel, Armando del Rio Hernandez, Charles Coombes, Giancarlo Pruneri, Adrian Lim, Fernando Calvo, Luca Magnani

## Abstract

Approximately 30% of women diagnosed with ERα breast cancer relapse with metastatic disease following adjuvant treatment with endocrine therapies^1,2^. The connection between acquisition of drug resistance and invasive potential is poorly understood. In this study, we demonstrate that the type II keratin topological associating domain (TAD)^3^ undergoes epigenetic reprogramming in cells that develop resistance to aromatase inhibitors (AI), leading to keratin 80 (KRT80) upregulation. In agreement, an increased number of KRT80-positive cells are observed at relapse *in vivo* while KRT80 expression associates with poor outcome using several clinical endpoints. KRT80 expression is driven by *de novo* enhancer activation by sterol regulatory element-binding protein 1^4^ (SREBP1). KRT80 upregulation directly promotes cytoskeletal rearrangements at the leading edge, increased focal adhesion maturation and cellular stiffening, which collectively promote cancer cell invasion. Shear-wave elasticity imaging of prospective patients shows that KRT80 levels correlate with stiffer tumors *in vivo*. Collectively, our data uncover an unpredicted and potentially targetable direct link between epigenetic and cytoskeletal reprogramming promoting cell invasion in response to chronic AI treatment.

## Main

AI treatment is standard of care for breast cancer (BC), yet BC cells frequently display drug-resistance and stronger metastatic potential at relapse, suggesting that chronic exposure to endocrine treatment might contribute in shaping the invasive potential, as suggested by previous *in vitro* studies^5^. The mechanism/s, order of events and molecular players mediating these phenomena are not well understood but it is likely that they involve cytoskeletal re-arrangements as they are essential for cancer invasion and metastasis ^6^. One possibility is that endocrine therapies (ET) might indirectly promote invasive potential by selecting for interrelated phenotypes during tumor evolution ^7–9^. Alternatively, AI treatment may directly contribute to the activation of invasive transcriptional programs. Chronic exposure to ET leads to coordinated activation and decommissioning of regulatory regions such as enhancer and promoters as shown by global changes in the localization of epigenetic marks H3K27ac and H3K4me1-2^5,9,10^. These epigenetic changes occasionally involve entire topological associating domains (TADs), three-dimensional compartments within the genome thought to restrict enhancer-promoter interactions ^3,11^. We observed that the type II keratin TAD ranked among the most significantly epigenetically reprogrammed (top 5%) when comparing untreated, non-invasive parental vs. invasive AI resistant BC cell lines^5^ (long term estrogen deprived: LTED cells, **Figure 1A**). Type I and Type II Keratins are the main constituents of cytoplasmic intermediate filaments and are involved in crucial cellular processes including cell attachment, stress adaptation and cell structure maintenance; yet very little is known about their role in cell movement and metastatic progression. Despite TAD dynamics, only few keratins within the type II-keratin TAD were transcriptionally reprogrammed in AI-resistant cell lines, including KRT80 (**Figure 1A and S1A-B**). Measuring KRT80 transcripts before or after short-term (48hrs) acute estrogen starvation using single cell RNA-seq data we showed that the significant increase in KRT80 positive cells is driven by transcriptional activation and not selection of KRT80-positive clones (**Figure 1B**). These data were validated in MCF7 and LTED cells using single cell RNA-FISH (**Figure 1C**). As expected, increased transcription corresponded to increased KRT80 protein level (**Figure 1D**). KRT80 is a largely unknown keratin structurally related to hair keratins^12^, in contrast with epithelial keratins commonly found in normal epithelial cells. This led us to further explore the role of KRT80 in promoting the aberrant phenotype observed in AI-resistant cells. KRT80 transcripts were also elevated in several ERα-negative cell lines, suggesting that upregulation in drug-resistant cells was not mediated by changes in ERα activity (**Table S1**). More importantly, IHC analysis of two independent clinical datasets confirmed that KRT80 positive cells increase after AI treatment but not Tamoxifen *in vivo*^13,14^ (**Figure 1E**). KRT80 localization *in vivo* was radically different to what has been shown in conventional keratins (e.g. KRT8, KRT14, KRT18 or KRT19^15^), presenting a peri-nuclear polarized pattern towards the lumen within healthy ducts and lobules (**Figure 1F and S2**). Similar staining patterns were conserved in benign lesions (**Figure S2**), whereas KRT80 staining became strongly cytoplasmic in higher grade BC and metastatic lesions suggesting a potential role in BC progression (**Figure 1E**). Correspondingly, high KRT80 mRNA levels correlated with poor survival in the TCGA-BC dataset (**Fig. 1G**). These data were confirmed by multivariate meta-analysis of two independent datasets with additional clinical endpoints (**Figure S3**).

**Figure 1.**
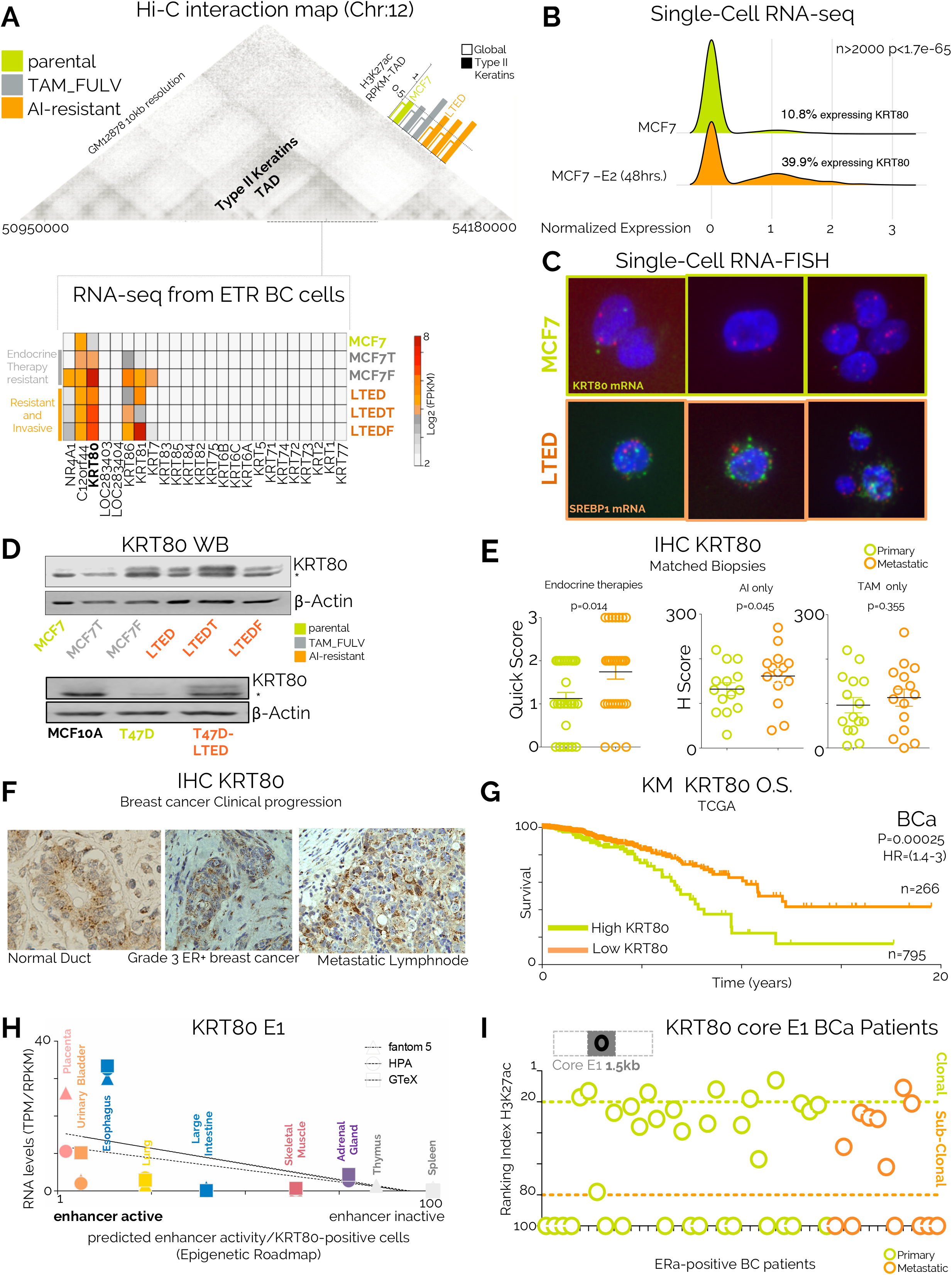
Keratin 80 transcription in AI-treated breast cancer cells is driven by enhancer activation. A) Hi-C 3D interactions in GM12878 cells were plot using http://promoter.bx.psu.edu/hi-c/view.php. The Type II keratin TAD boundaries were conserved in several other cell lines. The Type II TAD exhibits increased H3K27ac acetylation patterns in invasive drug resistant breast cancer cell lines compared to background (orange bars). The expression of Protein Coding genes within the TAD boundaries was assessed in parental treatment naïve (green), non-invasive drug resistant (grey) or invasive drug resistant breast cancer cell lines (orange) B) Single-cell RNA-seq data show that KRT80 increase is driven by transcriptional activation. Experiments were run comparing cells within 48 hours in absence of major cell division/apoptosis. Significance was calculated with a Fisher exact test C) Representative single-molecule, single cell RNA-FISH for SREBP1 (red) and KRT80 (green) confirm that LTED cells are characterized by a substantial increase in KRT80 transcription D) KRT80 protein levels are increased in two independent models of invasive drug resistant breast cancer cell lines when compared to parental or non-cancer breast cells. The asterisk represents an unspecific band. E) Matched clinical specimens from breast cancer patients show an increase in KRT80 positive cells following mono-treatment or sequential treatment with aromatase inhibitors. Similar results were not observed in Tamoxifen-only treated patients F) Immunocytochemistry (IHC) analyses show significant changes in KRT80 protein distribution between normal ductal cells (i), non-invasive (ii) and invasive ductal breast carcinoma cells (iii) E) KRT80 expression in diagnostic material is a bad prognosis marker G) Enhancer E1 activity is correlated with KRT80 RNA expression in several normal tissues H) Enhancer E1 activity is correlated with KRT80 RNA expression in several normal tissues using three independent databases (fantom5; HPA, Human Protein Atlas, GTEx, Genotype-Tissue Expression) I) Enhancer activity predicts the existence of dominant KRT80 subpopulations in primary and metastatic breast cancer patients.

Activation of cell type specific enhancers has been linked with cancer transcriptional aberration^9,16–18^, leading us to hypothesize that *de novo* enhancer activation within the TAD structure might control KRT80 expression in AI resistant cells. We used H3K27ac, an epigenetic mark associated with gene activation^9,19^, to identify KRT80 potential enhancers (**E1 and E2, Figure S4A**). As expected, E1-E2 activity was only captured in KRT80-positive cells (**Figure S4A**). 3D meta-analysis from parental MCF7 ChIA-Pet data strongly suggested that the E1 loci could contact the KRT80 promoter via enhancer-promoter interactions, while it excluded the weaker E2 (**Figure S4B**) suggesting that the 3D interaction is already pre-establihed in sensitive cells. To test whether E1 drove KRT80 transcriptional activity we adapted our recently developed computational pipeline to measure the relative size of KRT80-positive clones in several tissues^9^. Analysis of Epigenetic Roadmap data ^20^ strongly suggested that E1 activity controls KRT80 transcription levels (**Figure 1H**). E1 activity was also potentially associated with KRT80 transcription in several cell lines (**Figure S4C-D**). KRT80 E1 activity also correctly predicted strong expression in mammary epithelium cells (**Figure S4C-D**). Finally, E1 enhancer activity analysis predicted a significant increase in KRT80 positive cells in AI resistant models, in agreement with mRNA and protein analysis (**Figure S5**). Using fine-mapping analysis we identified a core-region within the E1 enhancer (1.5Kb) more strongly associated with KRT80 expression in our BC cell lines (**Figure. S5**). This core enhancer showed a clear pattern of activity in actual BC patients^9^ predicting the existence of KRT80 clonal and sub-clonal population in primary and metastatic BC (**Figure. 1I**). Overall, these data suggest that core-E1 is the critical enhancer driving KRT80 expression in BC cells.

We next investigated which transcription factor/s (TFs) might regulate KRT80 expression. DHS-seq analysis ^5^ indicated that KRT80 is already accessible in MCF7 (**Figure. 2A**), yet digital foot-printing suggested different occupancy rates (**Figure. 2B**). More specifically, we noted the appearance of a SREBP1 footprint within the core-E1 unique to LTED cells. We have previously reported that AI resistant cells upregulate lipid biosynthesis via global epigenetic reprogramming^5^ suggesting widespread SREBP1 activation in AI resistant cells. ENCODE TFs mapping showed that SREBP1 bind the core-E1 enhancers in lung cancer cells, the only ENCODE profiled cells characterize by strong KRT80 transcription (**Figure. S6A-B**). To directly test if SREBP1 drives KRT80 expression in BC we performed ChIP-seq in MCF7 and T47D cells and their respective AI-resistant models. Our data demonstrate that SREBP1 was bound at core-E1 only in AI-resistant BC cells (**Figure. 2C and S6C**). Interestingly, the expression of *KRT80* and SREBP1 target genes was also strongly correlated in BC patients (**Figure. S6D**). Finally, we show that SREBP1 silencing abrogated KRT80 expression in LTED cells (**Figure. 2D**). Overall these data demonstrate an unpredicted link between SREBP1 and KRT80 activation. Intriguingly, the footprint containing the SREBP1 motif was not under significant evolutionary constraint (**Figure. S7**), suggesting that the SREBP1-KRT80 axis might have evolved only recently.

**Figure 2.**
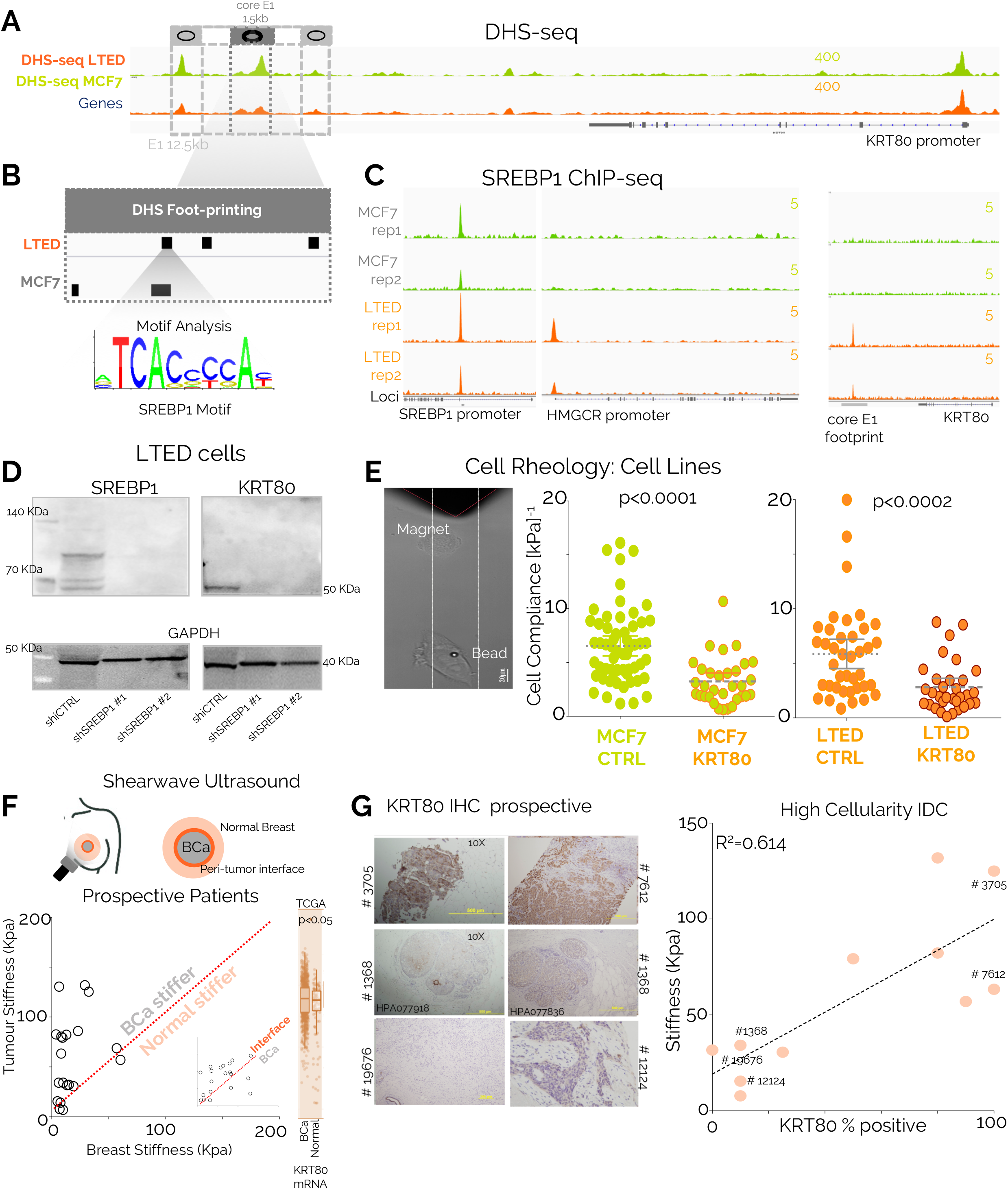
SREBP1 drives KRT80 expression leading to biomechanical changes in breast cancer cells. A) Open chromatin profiling via DHS-seq shows that the E1 enhancers is already accessible, but not yet acetylated in treatment naïve breast cancer cells (**Figure S4A**) B) Digital Foot-printing analysis suggest differential occupancy status within the E1 KRT80 enhancer C) ChIP-seq analysis demonstrate SREBP1 binding at the E1 core enhancers in invasive AI resistant breast cancer cells but not in treatment naïve parental cell lines D) SREBP1 depletion is sufficient to completely abrogate KRT80 protein levels in AI resistant cell lines E) Magnetic tweezer were used to measure the biomechanical properties of cells with or without KRT80 manipulation. KRT80 over-expression is sufficient to induce a surge in cellular stiffness in both parental MCF7 and AI-resistant LTED cells F) Shearwave Ultrasound measurement in prospective patients show that newly diagnosed breast cancer shows augmented stiffness compared to local healthy tissue. The cancer-stroma interface shows the highest increase in stiffness (inset) G) KRT80 cells in diagnostic material from prospective patients assessed with ultrasound were counted using IHC showing that intra-tumor stiffness correlates with the relative enrichment in KRT80 positive cells *in vivo*.

Several studies have investigated how mechanical stimuli influence the epigenetic landscape ^21,22^. However, our data implied a novel causal link whereby epigenetic reprogramming promoted changes in specific cytoskeletal components (e.g. KRT80) which may ultimately affect the biophysical properties of cells and tumors^23,24^ (**Figure 1–2**). In agreement, we observed a significant increase in cellular stiffness (inversely correlated to cell compliance) after KRT80 over-expression in MCF7 and LTED cells (**Figure 2E**). To test if KRT80 can contribute to tumor stiffness *in vivo* we prospectively recruited 20 patients with suspected BC and performed shear-wave elastography to measure intra-tumoral stiffness (**Figure 2F**). Our data showed that cancer lesions had significantly higher stiffness than surrounding normal tissues, with the highest peak of stiffness consistently measured at the invasive border (**Figure 2F**). Interestingly, meta-analysis of tumor and matched nearby tissue from TCGA show increased KRT80 mRNA in the tumor biopsies (**Figure 2F**). We then performed IHC for KRT80 with validated antibodies (**Figure 2G and S8**) using biopsies collected from our prospective patients. Linear regression analysis showed that KRT80 positivity significantly correlated with intra-tumor stiffness (**Figure 2G and S8**). Collectively, these data demonstrate that BCs characterized with high KRT80 content are mechanically stiffer.

The effect of increasing stiffness in metastatic invasion is highly debated. Previous studies have suggested that decreased stiffness, through loss of keratins, improves single-cell invasion^23^ typical of EMT cells. However, solid tumors can also use a myriad of multicellular invasion programs^25^ collectively termed “collective invasion”. Recent studies have shown that keratins such as KRT14 can play critical roles in collective invasion^26 6^ and multi-clonal metastatic seeding^26^, two processes driving BC progression^26^. In addition, a significant body of clinical literature has linked increased breast tumor stiffness to poorer prognosis ^26^ and lymph node positivity ^26^, independently of changes in extracellular matrix stiffness. We reasoned that a model in which KRT80 upregulation in BC cells leads to increased stiffness and augmented collective invasion might reconcile all these observations. To test this, we developed 3D spheroids from MCF7 or LTED cells and assessed collective invasion (**Figure 3A**) after KRT80 manipulation (**Figure S9A-D**). Spheroids from KRT80-positive LTED cells could effectively invade intro matrigel matrices, but KRT80 depletion completely abrogated the invasive phenotype (**Figure 3B-C and S9E-F**). Conversely, ectopic expression of KRT80 conferred matrix invading capacities in otherwise non-invasive MCF7 cells, even in the absence of chronic estrogen deprivation (**Figure 3B-C and S9E-F**). KRT80 immunostaining showed that KRT80 positive cells clustered at the invasive front in LTED spheroids (**Figure 3C**), a pattern reminiscent of the leading cells characterized in epithelial tumors during collective invasion^26^. Confocal microscopy analyses informed that LTED and MCF7-KRT80 cells presented an intricate network of KRT80 filaments that significantly overlap actin fibres. This KRT80 network was prominent in lamellipodium-like structures in leading cells (**Figure 3D-E and S10**). Conversely, in KRT80^low^ cells (i.e. MCF7 and LTED-sh^a^), KRT80 staining was punctuated and border cells presented strong cortical actin and no prominent lamellipodia^27^. Interestingly, KRT80 positivity strongly characterized cells from prospectively collected pleural effusion from AI treated patients (**Figure S10C**). Quantitative analysis of confocal data showed that KRT80 directly promoted the generation of actin fibers and larger more mature paxillin focal adhesions, two events associated with lamellipodia formation and cell stiffness/cellular tension^28,29^(**Figure 3E-3F and Fig. S10-11**).

**Figure 3.**
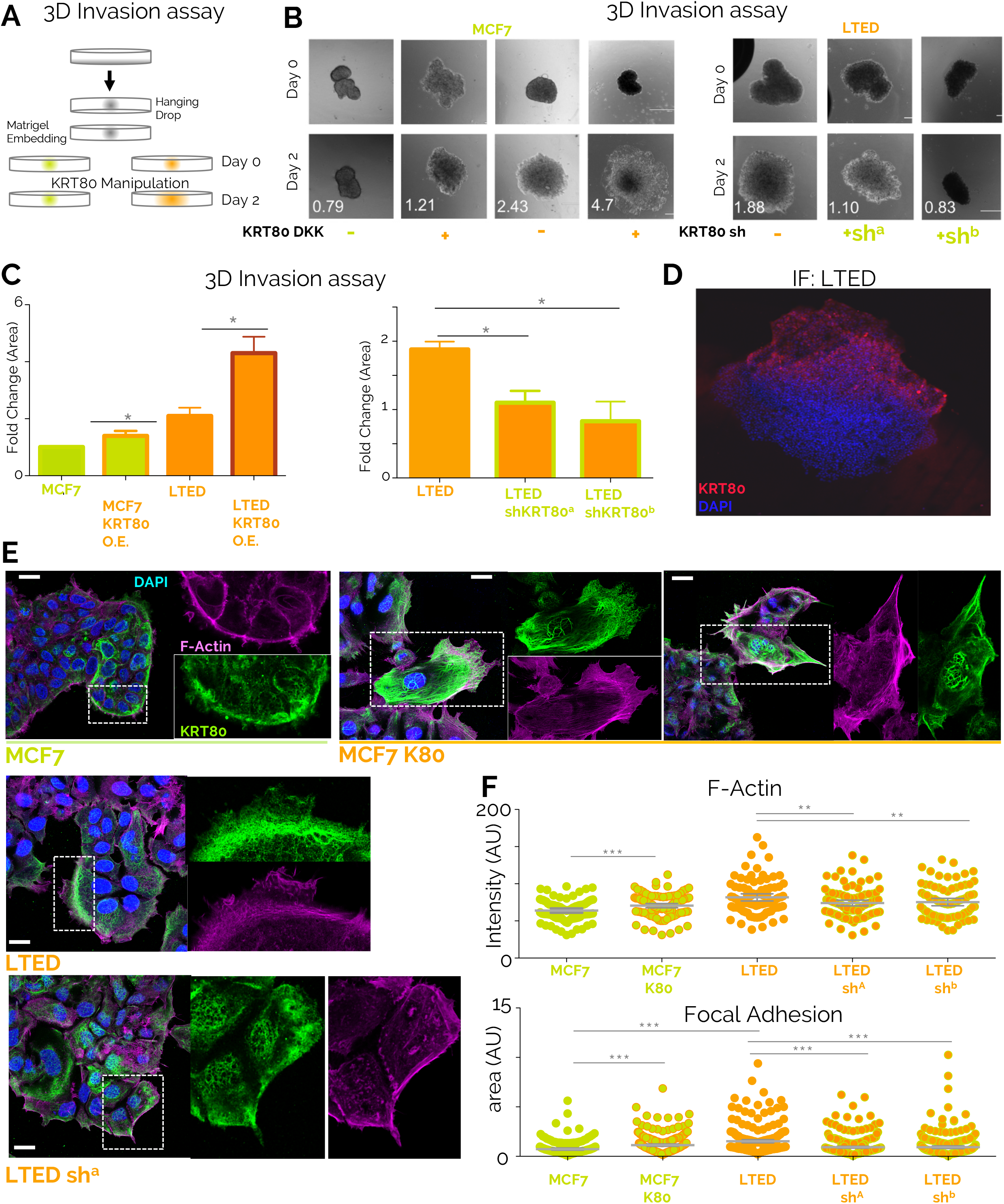
KRT80 increase directly promotes cell invasion. A) Design of the 3D invasion assay. Organoids were derived from treatment naïve (green; MCF7) or invasive AI resistant (orange; LTED) breast cancer cells. KRT80 expression was manipulated **via** ectopic overexpression or sh-mediated depletion. Organoids were embedded in Matrigel and monitored for 48 hours B) Representative brightfield images of KRT80-manipulated organoids show that increase of KRT80 further increases the invasive phenotype (i) whereas loss of KRT80 abrogates invasion (ii) C) Quantification of matrigel invasion assays shown in (B). D) Confocal microscopy of matrigel embedded invasive AI resistant LTED organoids E) E) Confocal microscopy show that KRT80 manipulation directly contribute to F-Actin accumulation at lamellopodia-like structures in breast cancer cell lines. Scale bars represent 25 μm. F) Quantification of F-Actin mean fluorescence intensity and Focal adhesion (pPaxilllin) area in function of KRT80 manipulation

To test if KRT80 manipulation drives ancillary phenotypes synergistic to cytoskeletal changes, we performed RNA-seq in cells transfected with KRT80 but where SREBP1 is not yet activated (non-invasive MCF7 cells, **Figure 4A**). Ectopic KRT80 expression led to clear transcriptional differences dominated by the reprogramming of a small set of genes (**Figure. 4A-B**). Pathway analyses of upregulated genes point out to cytoskeletal rearrangements (**Figure 4C**). Amongst them, we found particularly striking the strong KRT80-dependent induction of cortactin (CTTN), a factor directly linked to actin rearrangements, lamellipodia formation and cancer cell invasion^30,31^, that we confirmed by immunofluorescence (**Figure 4D**). In addition, we also detected a significant upregulation of *SEPT9*, a member of the septin family directly linked to actin fiber formation, focal adhesion maturation and motility ^32,33^ (**Figure 4B**). More importantly, genes activated in response to KRT80 activation have prognostic value, even when other classical clinical features are considered, in strong agreement with KRT80 prognostic features and suggesting that these genes might underlie early metastatic invasion (**Figure. 4E**). We also observed that several genes negatively regulated by KRT80 induction play central roles in cancer biology including negative regulators of migration (PCDH10, CADM1), tumor suppressors such as CDKN1A and PDCD2, genes involved in DNA repair (RAD50), chromatin remodelers as SMARCE1 and CHD4 and tumor specific antigens (CD276) suggesting a direct link between cytoskeletal reprogramming and several other oncogenic phenotypes (**Figure 4B**). Together, these results further support that KRT80 manipulation is sufficient to activate genes driving dramatic cytoskeletal rearrangements that ultimately induce invasive behaviors in BC and poorer prognosis. We cannot speculate at the moment if this is driven by a cytoskeleton-transcriptional feedback or is mediated by some specific transcriptional factors.

**Figure 4.**
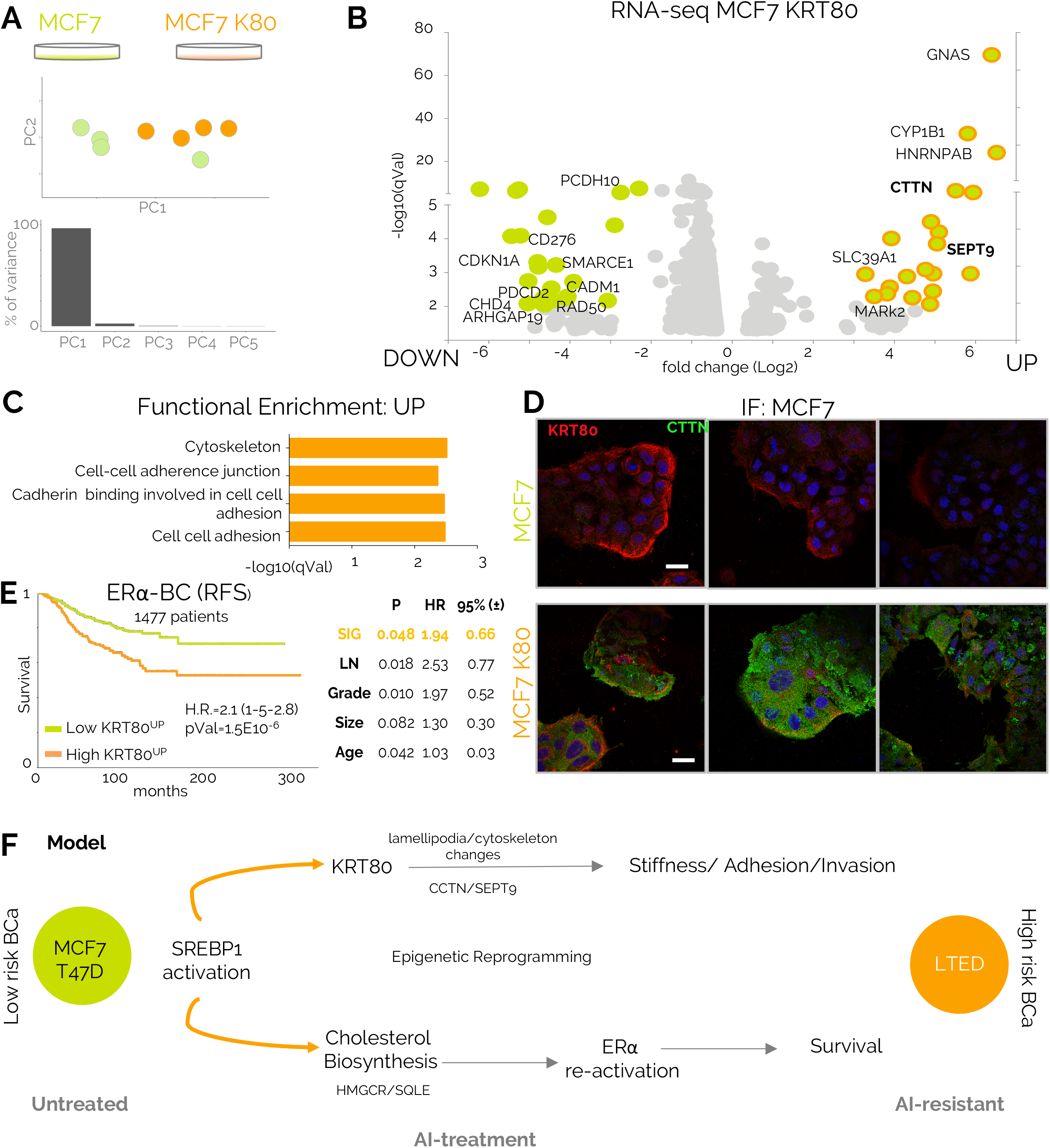
KRT80 increase promotes transcriptional changes of cytoskeletal genes. A) PCA analyses of RNA-seq profiled MCF7 breast cancer cells or MCF7 cells with ectopic expression of KRT80 B) Volcano plots of over-expressed or under-expressed genes in MCF7 cells following KRT80 ectopic expression C) Functional enrichment for gene upregulated following KRT80 ectopic expression D) Confocal imaging of Cortactin (*CTTN*) shows dramatic increase following KRT80 ectopic expression. Scale bars represent 25 μm E) Genes up-regulated in response to KRT80 over-expression (Figure 4B) have prognostic values in a multi-variate meta-analysis of microarray data obtained from 1477 ERα-positive breast cancer patients F) Current model: long-term AI treatment promotes constitutive activation of SREBP1 leading to pro-survival re-activation of estrogen receptor^5^ and global cytoskeletal re-arrangements. Cytoskeletal re-organization leads to direct biomechanics changes and promotes pro-invasive behaviour.

Aberrant cytoskeletal architecture characterizes tumor cells and is associated with cell migration and invasion; yet the mechanisms underlying cytoskeletal reorganization in tumour cells are not well understood. Here, we have uncovered a novel and causal link between endocrine therapy resistance, intra-tumoural stiffness and augmented invasive potential in luminal BC (**Figure. 4F**). Our data strongly suggest that therapy plays a direct role in shaping the biophysical properties and invasive potential of cancer cells, by inducing epigenetic rearrangements leading to KRT80 upregulation and concomitant cytoskeletal reorganization. We also describe an unexpected role for intermediate filaments in promoting cancer cell invasion by showing for the first time that KRT80 promotes actin fiber formation as well as focal adhesion maturation and lamellipodia formation. The link between epigenetic and cytoskeleton reprogramming offers an intriguing axis for drug development and biomarker discovery, especially within the goal of preventing metastatic invasion in BC patients treated with aromatase inhibitors.

## Acknowledgments

We want to acknowledge and thanks all patients and their families for the support and for donating the research samples. The authors gratefully acknowledge infrastructure support from the Cancer Research UK Imperial Centre, the Imperial Experimental Cancer Medicine Centre and the National Institute for Health Research Imperial Biomedical Research Centre. L.M was supported by a CRUK fellowship (C46704/A23110). Y.P was supported by a CRUK Studentship (PS2099). AJF and FC are funded by the Institute of Cancer Research, London (UK). FC is also funded by Worldwide Cancer Research (Grant 15-0273), Cancer Research UK (C57744/A22057) and the Ramon y Cajal Research Program (MINECO, RYC-2016-20352). We acknowledge Z. Magnani for his constructive comments on the manuscript.

## Author Contribution

**Competing Financial Interests Statement**

The authors declare no competing interests.

## Online Materials and Methods

### Cell lines and cell culture

In this study we used MCF7 breast adenocarcinoma cell line and derived resistant clones (Supplementary Figure 1). MCF7 Tamoxifen Resistant cell line (MCF7TR) was derived from MCF7 upon one year treatment with Tamoxifen. LTED (Long Term Estrogen Deprivation) cell lines were derived from MCF7 cell line upon one year estrogen deprivation, mimicking aromatase inhibitor resistance. LTED Tamoxifen Resistant cells (LTEDTR) were derived from LTEDs upon one year Tamoxifen treatment. Additionally, we employed an alternative aromatase inhibitor resistant model: T47D breast adenocarcinoma cell line and T47D-LTED. The latter was derived from T47D parental upon six months of estrogen deprivation. MCF7 and T47D breast carcer cell lines were cultured in DMEM (Sigma-Aldrich) supplemented with 10% FCS (Foetal Calf Serum, First Link UK), 2 mM L-Glutamine, 100 units/mL penicillin, and 0.1 mg/mL streptomycin (Sigma). MCF7 were further supplemented with 10-8 M Estradiol (Sigma). Estrogen-deprived cell lines (LTED, LTEDTR and T47D-LTED) were cultured in phenol-red free DMEM (Gibco, Life Technologies) supplemented with 10% DC-FCS (Double Charcoal stripped Foetal Calf Serum, First Link UK) and 2 mM L-Glutamine, 100 units/mL penicillin, and 0.1 mg/mL streptomycin (Sigma-Aldrich). LTEDTR were further supplemented with 10-7 M Tamoxifen (SIGMA).

### Generation of stable cell lines

For KRT80 overexpression, a full length KRT80 cDNA clone Myc-DKK-tagged was obtained from OriGene and transformed into DH5α competent cells (Invitrogen). Plasmidic DNA was isolated using Maxi-Prep Kit (QIAGEN) and transfected in MCF7 and LTED cells using X-tremeGENE 9 DNA Transfection Reagent (Roche) following manufacturer’s instructions. Transfected cells, carrying Neomycin resistance, were selected with G418 (SIGMA), used at a final concentration of 1 mg/mL for MCF7 and 0.5 mg/mL for LTED. Knock-down of KRT80 was achieved by transfection of two different shRNA expression vectors and a scrambled negative control obtained from OriGene. Cells carrying the corresponding construct were selected with Puromycin (Sigma-Aldrich) at a final concentration of 1 ug/mL for MCF7 and 0.5 ug/mL for LTED cell line.

### RNA extraction and RT-qPCR

Cells were washed with PBS and harvested using a cell lifter (Corning) in RLT buffer supplemented with 1% β-mercaptoethanol. Cell lysate was homogenized using QIAshredder columns (QIAGEN) and RNA extraction was performed with RNeasy Mini Kit (QIAGEN) following manufacturer’s instructions. RNA concentration was measured using a NanoDrop 1000 Spectrophotometer and 0.5-2 μg of RNA were retrotranscribed using High Capacity cDNA Reverse Transcription Kit (Applied Biosystems). Quantitative PCR (qPCR) was performed using 2X SYBR GREEN Mix (Invitrogen) and expression levels of each gene were calculated using the 2-ΔΔCt method, normalizing expression levels to 28S transcript.

### Protein extraction, quantification and western blotting

Cells were harvested in 50μL ice-cold RIPA buffer (50 mM Tris-HCl at pH 8.0, with 150 mM sodium chloride, 1.0% Igepal CA-630 (NP-40), 0.5% sodium deoxycholate, and 0.1% sodium dodecyl sulphate) (Sigma; #R02780), supplemented with 1X protease (Roche; #11697498001) and 1X phosphatase (Sigma; #93482) inhibitor cocktail. The cell pellet and RIPA were mixed by pipetting up and down, incubated at 4°C for 30 minutes and vortexed every 5 minutes. Cell lysates were then centrifuged at 13,000 rpm for 30 minutes at 4°C. The supernatants were transferred to a new 1.5mL eppendorf tube and the pellets were discarded. Protein concentration was measured using BCA Assay Kit (Thermo Fisher) following manufacturer’s instructions. With regard to western blotting, 20μg of protein per sample, were mixed with 4X Bolt sample buffer (Life Technologies; #B0007), 10X Bolt sample reducing agent (Life Technologies; #B0009), ddH2O and heated at 95°C prior to loading. Protein lysate were loaded into BOLT 4-12% Bis-Tris Plus Gel (Life Technologies; NW04120BOX). The pre-made gel was placed into a mini gel tank (Life Technologies; #A25977) containing 1X Bolt running buffer (Life Technologies). Electrophoresis was carried out at 90V for 35 minutes to allow proteins to adequately run through and also until the bromophenol blue dye reached the bottom of the gels. The gels were transferred into a Biotrace nitrocellulose membrane (VWR; #PN66485) using a TE-22 transfer unit (Hoefer GE Healthcare) at 100V for 90 minutes. The membrane was incubated in blocking buffer for 45 minutes at room temperature to reduce non-specific binding of primary antibody. The membrane was then incubated with the diluted primary antibodies (Anti-KRT80 HPA 077836 Atlas Antibodies (1:200 dilution), Anti-SREBP1 H-160 sc-8984 Santa Cruz Biotechnology (1:200 dilution), (Guinea Pig Anti-KRT80 1:5,000; Mouse Anti-DKK 1:1,000, OriGene; Mouse Anti-β-Actin 1:10,000) in blocking buffer at 4°C and allowed to shake overnight. After primary antibody incubation, the membrane was washed three times in PBST (5 minutes per wash on a rocking platform) and then incubated for 1 hour with the HRP-GAPDH (Abcam; #ab9482 (1:5000 dilution)) conjugated antibody (for the loading control membrane) which was diluted in 5% BSA/PBST and goat anti-rabbit IgG (H+L) Cross Absorbed secondary antibody, HRP 1:20000 dilution (ThermoFisher Scientific; #31462). The membranes (including the loading control membrane) were washed three times in PBST. Amersham ECL start Western Blotting Detection reagent (GE Healthcare Life Sciences; #RPN3243) was used for chemiluminescent imaging using the Fusion solo (Vilber; Germany) imager.

### Chromatin Immunoprecipitation (ChIP)

For ChIP, cells were fixed with 1% formaldehyde for 10 min at 37°C and reaction was quenched with 0.1M glycine. The cells were subsequently washed twice with PBS after which they were lysed in lysis buffer (LB) 1 (50 mM HEPES-KOH, pH 7.5, 40 mM NaCL, 1mM EDTA, 10% glycerol, 0.5% NP-40, 0.15% Triton X-100), for 10 minutes, then for 5 minutes in LB 2 (10 mM Tris-HCl, pH 8.0, 200 mM NaCl, 1 mM EDTA and 0.5 mM EGTA) and subsequently eluted in LB 3 for sonication (10 mM TRIS-HCl, pH 8.0, 100mM NaCl, 1 mM EDTA, 0.5 mM EGTA, 0.1% Na-Deoxycholate and 0.5%N-lauroylsarcosine). DNA was sheared using the Bioruptor^®^ Pico sonication device (High, 10 cycles of 30’’ on and 30’’ off) (Diagenode). Sheared chromatin was cleared by centrifugation. Magnetic beads were precoated by adding 10 μg of antibody Rabbit-anti-SREBP1 (H-160): sc-8984 (Santa Cruz Biotechnology, Inc.) to 50 μl magnetic beads per ChIP (Dynabeads protein A, Life technologies) and incubated for 6 hours on a rotating platform at 4°C. Diluted sheared chromatin was added to the coated magnetic beads and incubated on a rotating platform at 4°C O/N. 10 μl of sheared chromatin taken as input and treated the same. The next day magnetic bead complexes were washed three times with RIPA buffer (50 mM HEPES pH 7.6, 1mM EDTA, 0.7% Na deoxycholate, 1% NP-40, 0.5 M LiCL) and two times with TE buffer (10mM Tris pH 8.0, 1 mM EDTA). DNA is O/N eluted from the beads in 100 μl de crosslinking buffer (50 mM Tris-HCl, pH 8.0, 10 mM EDTA, 1% SDS) at 65 °C. After overnight de-crosslinking, 200μl TE buffer was added and the eluted DNA was treated with 8 μl of 1mg/ml RibonucleaseA (RNaseA) for 30 min at 37°C and subsequently incubated with 4 μl of 20 mg/ml proteinase K (Invitrogen) for 1 hour at 55 °C. Then DNA extraction was performed using SPRI magnetic beads (Beckman Coulter, B23318). After elution in TE buffer, DNA was quantified using Qubit (ThermoFisher Scientific; Qubit 3.0 Fluorometer; #Q33216) high sensitivity assay (ThermoFisher Scientific; #33216). Quantitative polymerase chain reaction (qPCR) was then carried out (Applied Biosystems; #7900HT Real time PCR, #StePOnePlus). If sufficient enrichment is seen in the antibody treatment samples over the ‘input’ samples and compared to internal negative controls, these undergo DNA size selection and library preparation.

### Library preparation

Prior to sequencing, ChIP samples were library prepared using the NEBNext Ultra II DNA Library Prep Kit for Illumina (New England Biolabs, NEBNext Ultra II DNA library prep kit for Illumina, #E7770, NEBNext Multiplex Oligos for Illumina, #E7335L). Adaptor ligated DNA is size selected with SPRI magnetic beads (Beckman Coulter, B23318) which aims to retain DNA fragments between 200-300 base pairs (bp), recognisable for the Illumina sequencer (#NextSeq500). After library preparation, we performed qPCR, high sensitivity DNA quantification and size selection measurement (Agilent Bioanalyzer 2100 system + High sensitivity DNA measurement assay; 5067-4626) before sending samples for sequencing.

### RNA sequencing

Total RNA from each sample was quantified by Qubit^®^ Fluorometer and quality checked by Agilent Bioanalyzer^®^ RNA 6000 Nano Chip. All samples have high quality RNA with a RIN score > 7. One microgram of total RNA from each sample was used as starting material for paired-end RNA-seq library preparation using NEBNext rRNA Depletion Kit (NEB #E6310) and NEBNext Ultra II RNA Library Prep Kit for Illumina (NEB #E7770) following the manufacturer’s instructions. Libraries were sequenced on an Illumina Next Seq machine (#NextSeq500). Reads were processed using Kallisto and DEGS were called using Sleuth ^34^

### Single cell RNA-FISH

Cell were cultured, fixed and pretreated according to the protocol for the RNAscope^®^ Multiplex Fluorescent Reagent Kit v2 Assay provided by Advanced Cell Diagnostics (ACD, #323100, Nunc Lab Tek II 2 Well Glass Slides, #154461K). The assay was run following the manufacturer’s instructions, hybridization was performed overnight. PerkinElmer TSA Plus Fluorophores (fluorescein, NEL741001KT and Cyanin 3, NEL744001KT) were diluted at 1:1300 and assigned to the channels HRP-C1 and HRP-C2, respectively.

| Fluorophores | Part number |
| --- | --- |
| PerkinElmer TSA Plus Fluorescein System | NEL741001KT |
| PerkinElmer TSA Plus Cyanine 3 System | NEL744001KT |

Samples were imaged using a ×60 objective with a Ti Nikon microscope equipped with a spinning disk (CAIRN) and analysed in Image J.

| Catalog Number | Product Name | Probe Type | Channel | Species Name |
| --- | --- | --- | --- | --- |
| 320861 | RNAscope® 3-plex Positive Control Probe | RNAscope® Control Probes | 2 | Homo sapiens |
| 310043 | RNAscope® Negative Control Probe - DapB | RNAscope® Control Probes | 1 | Other* |
| 300031 | RNAscope® Target Probe C1 | RNAscope® 20ZZ probe named Hs-KRT80 targeting 294-1445 of NM_182507.2 | 1 | Homo sapiens |
| 300031 | RNAscope® Target Probe C2 | RNAscope® RNAscope® Probe - Hs-SQLE 465071 | 2 | Homo sapiens |
| 300031 | RNAscope® Target Probe C2 | RNAscope® 20ZZ probe named Hs-SREBF1 targeting 958-2002 of NM_001005291.2 | 2 | Homo sapiens |
| 470561 | RNAscope® Probe - Hs-HMGCR | RNAscope® Target Probes | 1 | Homo sapiens |

\**Bacillus subtilis* dihydrodipicolinate reductase (dapB) gene

### 3D Organoid assay

250,000 cells were resuspended in 1 mL of the corresponding media and 20 μL drops were placed in the lid of a 10 cm dish (Corning). The lid was flipped over the dish containing 5 mL of media in order to prevent evaporation. Hanging drops were incubated for 5 days at 37% C in a humidified atmosphere, during which formation of organoids was achieved. To follow, organoids were immersed in 10 μL of phenol-red free Matrigel^®^ (BD Biosciences) and placed in a 24 well-plate (Corning) The appropriate media containing G418 or puromycin was subsequently added to the well. Brightfield images were acquired at days zero and day two using an EVOS microscope (Advanced Microscopy Group, Life Technologies). Images were analysed using Fiji ImageJ software and fold-change area was calculated using the following formula: Area (fold-change) = Area Day 2/Area Day 0

### Immunofluorescence and confocal microscopy

Organoids were washed with PBS and fixed for 15 minutes with 4 % PFA/PBS. Fixation was stopped by rinsing with 100 mM Glycine/PBS. Cells were permeabilized with 0.5% Triton/PBS X-100 and unspecific binding was blocked with blocking solution (5% BSA, 0.2 % Triton X-100, 0.05% Tween in PBS) for 90 minutes. Organoids were then incubated with primary antibody (Rabbit Anti-KRT80 1:200, Sigma-Aldrich) for 2 hours, washed three times with washing buffer (0.2 % Triton X-100, 0.1 % BSA, 0.05% Tween in PBS), and incubated with secondary antibody (Goat Anti-Rabbit Alexa Fluor 555 1:200, Invitrogen) for 45 minutes. Organoids were washed with immunofluorescence buffer for 20 minutes and PBS for 10 minutes. Finally, organoids were mounted in Moviol (AppliChem) containing 5 μg/mL of DAPI (Lonza) and visualized using a Zeiss LSM-780 inverted confocal microscope.

### Immunofluorescence and Image Analysis

Cells were seeded on glass bottom 24 well plates (MatTek) coated with 10 μg/ml fibronectin (Sigma), fixed in 4% PFA and permeabilized in PBS with 0.2% Triton X. The samples were blocked in 3% BSA with 0.1% PBS Tween (PBST) for 3 hours. The primary antibodies (Cortactin, 05-180, Millipore, 1:100; Keratin-80, HPA077836, Sigma Atlas, 1:100; Phospho-Paxillin-pY118, 44-722g, Invitrogen, 1:100) were diluted in 3% BSA in 0.1% PBS Tween and incubated overnight at 4°C. The wells were then washed 3 times in 3% BSA 0.1% PBST for 10 min, followed by the addition of the appropriate secondary antibody (Alexa Fluor, Invitrogen), DAPI (Sigma) and FITC-phalloidin (Sigma). Imaging was performed using Leica SP8 Confocal microscope. Image analysis was performed using Volocity (Perkin Elmer), evaluating the mean F-Actin fluorescence intensity of approximately 100 cells per cell line and the area of approximately 250 focal adhesions per cell line.

### Tissue specimens

Seventy-five human breast specimens and ten metastatic lymph nodes were selected from Histopathology Department at Charing Cross Hospital, with the previous approval of Imperial College Healthcare NHS Trust Tissue Bank.

A Tissue Microarray (TMA) containing 26 primary breast tumors and paired ETR relapses was constructed as previously described (*18*).

Immunohistochemistry staining was scored using a quick score system by two independent investigators, one of them a consultant pathologist (SS). Score was calculated as follows: S=3 (strongly stained cells), S=2 (moderate staining), S=1 (poorly stained cells) and S=0 (absence of staining). Staining intensity was assessed as mean intensity from the tumor region contained within the TMA.

### Immunohistochemistry

Formalin fixed and paraffin embedded (FFPE) tissue specimens were sliced in 4 μm sections using a Leica RM2235 manual microtome. Dried sections were de-waxed by immersion in xylene and rehydrated with subsequent immersion in 100% ethanol, 70 % ethanol and distilled water. Antigen retrival was performed by immersion in PBS 0.01 M citric acid pH 6 and heated at 800 W during 15 minutes. Slides were rinsed in PBS and endogenous peroxidase activity was blocked during 30 minutes using Dako RealTM Peroxidase Blocking Solution. Following that, slides were rinsed twice with PBS and incubated with 10% pig serum (Bio-Rad) during 30 minutes and overnight with KRT80 antibody (Sigma-Aldrich, 1:200). Following day, slides were rinsed in PBS and incubated 30 minutes with secondary antibody (biotinylated Goat Anti-Rabbit IgG 1:200, Vector Laboratories) and 30 minutes with an avidin/biotin peroxidase-based system (VECTASTAIN Elite ABC Kit, Vector Laboratories). Colour reaction was developed during 1 minute using DAB (Diaminobenzidine, Vector ImmPACT DAB Peroxidase Substrate). Colour development was stopped by immersion during 5 minutes in running tap water and following that, nuclei was stained with haematoxylin. Slides were dehydratated in 100% ethanol, cleared in xylene and mounted in DPX (SIGMA).

### Statistical analysis

Data is presented as mean ± SD (standard deviation). Data analysis was performed using GraphPad Prism 6 software. An unpaired two-tailed Student’s t test was applied to all data, except from a non-parametric Mann-Whitney test applied to TMA score.

### Survival analysis

Publicly available breast cancer datasets were identified in GEO (https://www.ncbi.nlm.nih.gov/geo/), EGA (https://www.ebi.ac.uk/ega/home), and TCGA (https://cancergenome.nih.gov/). Only cohorts including at least 30 patients and with available follow-up data were included. Samples derived using different technological platforms (Affymetrix gene chips, Illumina gene chips, RNA-seq) were processed independently. For KRT80, the probe set 231849_at was used in the Affymetrix dataset, the probe ILMN_1705814 was used in the Illumina dataset and the gene 144501 was used in the RNA-seq dataset. Cox proportional hazards survival analysis was performed as described previously^35^. Kaplan-Meier plots were derived to visualize survival differences. In the multivariate analysis, the RNA expression of ERα, HER2, and MKI67 were used as surrogate markers for ER and HER2 status, and for proliferation. In this, the probe sets 205225_at, 216836_s_at, and 212021_s_at were used for ERα, HER2, and MKI67, respectively. The survival analysis was performed for relapse-free survival (RFS), overall survival (OS), and post-progression survival (PPS). PPS was computed by extracting the RFS time from the OS time for patients having both RFS and OS data and having an event for RFS. Censoring data for PPS was derived from the OS event. The survival analysis was performed in the R statistical environment.

### Cellular microrheology

To characterize the mechanical properties of the four different BC cell lines, we used magnetic tweezer microrheology to measure cell deformation in response to magnetically generated forces. Tensional magnetic forces were induced by a high gradient magnetic field generated by an electromagnetic tweezer device. The positioning of the tip of the magnetic tweezer device was controlled by an electronic micromanipulator. Superparamagnetic 4.5 μm epoxylated beads (Dynabeads, Life Technologies) were coated with fibronectin (40 μg per 8 x 10^7^ beads, Sigma Aldrich F0895) and incubated with adherent cells for 30 minutes, prior to measurements, to allow integrin binding and provide a mechanical link between the bead and the cytoskeleton. The unbound beads were removed by multiple washing with PBS. The experiments were performed at 37°C, 5% CO2 and 95% humidity in DMEM containing 2% FBS in a microscope stage incubation chamber. A viscoelastic creep experiment was conducted by applying mechanical tension onto single beads bound on the apical surface of the cells with a constant pulling force (F_0_ = 1 nN) for 3 seconds generated by the magnetic tweezers. The viscoelastic creep response of the cells was recorded by tracking the resulting bead displacement in brightfield (40x objective at 20 frames per second, Nikon Eclipse Ti-B) that is indicative of the local cytoskeletal deformation. A custom-built MATLAB algorithm was then used to analyse the image sequences and track bead displacement by following the intensity-weighted centroid of the bead across all captured frames. The viscoelastic creep response J(t) of cells during force application followed a power-law in time J(t) = J_0_(t/t_0_)^β^ with the prefactor J0 representing cell compliance (J_0_ = inverse of cell stiffness in units of kPa^-1^) and the dimensionless exponent β representing cell fluidity with values ranging between 0<β<1 pure elastic (β = 0) or viscous behaviour (β = 1) and with the reference time t0 was set to 1 sec. The creep compliance J(t) represents the ratio ( γ(t)/σ_0_) of the localized cellular strain γ(t) induced by the applied stress from the magnetic tweezers σø, with γ(t) taken as the radial bead displacement normalised over the bead radius γ(t)=d(t)/r and the applied stress as σ_0_ =F_0_/4πr^2^ taken as the applied force normalised over the bead cross sectional area. Compliance measurements for each BC cell line were collected from three independent experiments (MCF7 CTRL n=60, MCF7 KRT80 n=34, LTED CTRL n=41, LTED KRT80 n=34).

### Shearwave Elastography

All SWE was performed by a breast radiologist with more than 10-years experience of performing Breast ultrasound and elastography on breast lesions. A state-of-the art ultrasound scanner, Aplio i900 (Canon Medical Systems, Nasu, Japan) with the latest 2D SWE technology was used for this study. All SWE maps and calculations were obtained pre-biopsy. A good stand-off was used for superficial lesions and initially, continuous SWE mode ("multi-shot") was used to select the optimum plane and once this was stabilised, a higher energy SWE push-pulse ("one-shot" mode) was then utilised to obtain the final elastogram for calculations. Regions of interest (ROI) were placed within the centre of the lesion, in the periphery and also within the adjacent normal breast tissue. This has been stored as raw data within the ultrasound systems which would enable any re-calculations as necessary.

## Supplementary Figures

**Supplementary Figure 1: KRT80 expression in BC cell lines**. A) Genes that were found differentially regulated in RNA-seq analysis were investigated using qRT-PCR in durg resistant cell lines derived from drug sensitive MCF7 cells B) KRT80 transcripts were also quantified in non-tumorigenic breast MCF10A cell lines and in drug sensitive and drug-resistant clones of T47D breast cancer cell lines. Bars and error bars represent the average and the SD of 3 biological replicates. One-Way ANOVA with Dunnet’s correction was used to establish statistical significance. Asterisks represent significance levels at *, **, *** P<0.05, 0.01 and 0.001 respectively

**Supplementary Figure 2: KRT80 IHC analysis in normal breast and various breast cancer pathologies**. A-D) KRT80 was imaged using IHC in a series of human samples collected at Charing Cross Hospital (London). Tissues were collected to cover a large spectrum of benign and malignant lesions including metastatic samples from Breast Cancer patients.

**Supplementary Figure 3: KRT80 mRNA levels have prognostic significance**. A) KRT80 mRNA levels were used to stratify breast cancer patients profiled using microarray technologies. All histologies were included in the analysis B) Multivariate correction analysis for panel A. High levels of KRT80 mRNA remains significantly associated with faster relapse even when other significant clinical parameters are accounted for. The three bottom panels show KRT80 prognostic significance in the context of post-progression survival (PPS), distal-metastasis free survival (DMFS) and overall survival (OS) C) KRT80 is associated with poor overall survival in a large independent cohort of ERα breast cancer patients profiled with Illumina Microarray technologies D) KRT80 prognostic significance was investigated in three subcohorts derived from the METABRIC cohort and were tested in function of prognostic power in the short-term (<5 years) or long-term (<25years). Hazard ratios are plotted on the x-axis (>1= worse overall survival) E) KRT80 prognostic association with overall survival was tested in patients which had annotated post-surgical adjuvant treatment.

**Supplementary Figure 4: KRT80 enhancer mapping**. A) H3K27ac ChIP-seq track for MCF7 and associated drug-resistant cell lines in the Type II kerating TAD locus identifies E1 and E2 as two potential enhancers involved in KRT80 regulation B) Virtual 4C (http://promoter.bx.psu.edu/hi-c/virtual4c.php) using POL2 Chia-PET data to predict potential loops between the E1 and E2 loci and KRT80 promoter in MCF7 cells C) Enhancer clonality analysis of ENCODE profiled cell lines predict E1 as being clonal in KRT80 positive cell lines D) RNA levels for KRT80 in a large panel of cell lines identifies KRT80 positive cells in few tissues in which E1 appears to be clonal.

**Supplementary Figure 5. E1 fine mapping**. Enhancer clonality analysis reveal that E1 contains sub-enhancers with various degree of predicted clonality in MCF7 and associated drug-resistant cell lines. Enhancer clonality drastically increase in KRT80 positive cells suggesting that E1 controls KRT80 expression.

**Supplementary Figure 6. E1 fine mapping**. A) E1 contains three major sub-enhancers as shown by DNAseI-hypersensitive mapping in several cell lines. Lung cancer A549 also shows potential activation of E1 enhancers B) Within the ENCODE cohort, only A549 lung cancer cell lines show KRT80 expression C) ChIP-seq analysis of SREBP1 genome-binding demonstrate that SREBP1 binds the E1 enhancer in KRT80 positive T47D LTED cells but not in KRT80 negative T47D cells D) Metanalysis of highly correlated RNA transcripts shows that KRT80 and classical SREBP1 targets show strong correlation in clinical samples.

**Supplementary Figure 7. E1 is not conserved**. Conservation analysis in the E1 locus and nearby area shows that E1 sequence containing the SREBP1 motif has emerged relatively late in evolution and is not found in other mammalian species.

**Supplementary Figure 8. Additional validation of KRT80 antibodies**. We have conducted additional analysis to validate two independent antibodies for KRT80 in breast tissues and breast cancer samples. Antibody are labelled and were obtained from the Protein Atlas Initiative

**Supplementary Figure 9. Functional characterization of KRT80**. A) RNA expression analysis shows that over-expression of exogenous KRT80 induce increased expression in MCF7 and LTED cells B) WB analysis using KRT80 or DKK antibodies confirmed KRT80 upregulation upon exogenous expression of the KRT80 construct C) Stable transfection of two independent shRNAs targeting KRT80 can reduce KRT80 expression and protein levels D) Transient transfection of two independent siRNA targeting KRT80 can reduce KRT80 expression and protein levels E) Migration analysis of transiently transfected siRNA-KRT80 cells show significant reduction in migratory capacity F) Matrigel invasion analysis of transiently transfected siRNA-KRT80 cells show significant reduction in invasiveness capacity. Invasion and Migration experiment using transient siRNA were conducted in biological triplicates using invasive and migratory LTED cell in which KRT80 is highly expressed. Bars and error bars represent the average and the SD of 3 biological replicates. One-Way ANOVA with Dunnet’s correction was used to establish statistical significance. Asterisks represent significance levels at *, **, *** P<0.05, 0.01 and 0.001 respectively

**Supplementary Figure 10. KRT80 is preferentially distributed at the margin of 3D cultures**. A) IF analysis of KRT80 in MCF7 cells and drug-resistant derivatives. KRT80 appears to be enriched in filopodia-like structures B) IF imaging shows that when cells are grown as organoids, KRT80 positive cells accumulate at the matrigel-organoid border. Blown up show that KRT80 appears to be enriched in filopodia like structures actively penetrating the matrigel structure C) IF analysis shows that KRT80 is strongly expressed in cells cancer cells isolated from pleural effusion from breast cancer patients

**Supplementary Figure 11. KRT80 enriched substructures are occupied by F-Actin and focal adhesion molecules**. High-resolution confocal analysis shows that KRT80 enriched filipodia-like structures contain F-Actin and focal adhesion molecules. Manipulation of KRT80 directly reprogram these sub-structures.

